# Microgroove substrates unveil topography-driven, dynamic 3D nuclear deformations

**DOI:** 10.1101/2024.02.02.578638

**Authors:** Claire Leclech, Bettina Roellinger, Joni Frederick, Kamel Mamchaoui, Catherine Coirault, Abdul I. Barakat

## Abstract

Navigating complex extracellular environments requires extensive deformation of cells and their nuclei. Nuclear deformations are intricately linked to nuclear structure and mechanical properties, and abnormalities in nuclear mechanics contribute to various diseases including laminopathies and cancer. Most *in vitro* systems used to study nuclear deformations are typically designed to generate strong whole-cell confinement relevant for specific cell types such as immune or cancer cells. Here, we use microgroove substrates as a model of anisotropic basement membrane topography and we report that adherent cells including endothelial cells and myoblasts exhibit significant 3D (in-plane and out-of-plane) nuclear deformations, with partial to complete penetration into the microgrooves. These deformations are dynamic with nuclei cyclically entering and exiting the microgrooves. AFM measurements show that these deformation cycles are accompanied by transient changes in nuclear mechanical properties. We also show that nuclear penetration into the grooves is principally driven by cell-substrate adhesion, without the need for cytoskeleton-associated forces. Finally, we demonstrate that myoblasts from patients with *LMNA* mutations exhibit abnormal nuclear deformations which can be rapidly identified and quantified using automated image analysis. We therefore propose the use of microgrooves as a novel simple, tunable, and high throughput system to study nuclear deformations in adherent cells, with the potential to serve as a functional diagnostic platform for pathological alterations in nuclear mechanics.

## Introduction

Cells *in vivo* are often embedded in intricate environments whose geometrical organization simultaneously imposes physical guidance and mechanical confinement. Navigating these complex spaces requires extensive deformation of the nucleus, the largest organelle in the cell and a major cellular mechanostransducer (Dahl *et al*, 2008; Kirby & Lammerding, 2018). The structural integrity of the nucleus relies principally on nuclear envelope proteins, most notably lamins, that provide a scaffold that safeguards chromatin organization, thereby ensuring genome integrity in the face of mechanical stress (Stephens *et al*, 2017; Stephens, 2020; Hobson *et al*, 2020). It is therefore not surprising that altered nuclear mechanical properties are observed in a number of pathologies including laminopathies and cancer (Zwerger *et al*, 2011). In laminopathies, mutations in genes encoding A-type lamins can lead to severe disorders such as cardiomyopathy, muscular dystrophy, or premature aging (progeria) (Hah & Kim, 2019; Worman, 2012). In the case of cancer, abnormal nuclear morphology is often a hallmark of the disease, and increased nuclear deformability correlates with heightened tumoral invasion (Denais & Lammerding, 2014).

Different *in vitro* tools have been developed over the years to study nuclear rheology and deformations in various cell types (Hobson *et al*, 2021). One class of such systems, which includes atomic force microscopy (AFM), optical/magnetic tweezers, and micropipette aspiration, enables the application of controlled forces that lead to local stretching or compression of individual nuclei, thereby providing precise mechanical characterization. These systems, however, suffer from a number of limitations including high experimental complexity, low throughput, and limited direct physiological relevance. Other systems, such as microfluidic channels with constrictions, constrain the cells uniformly over their entire contact surface and are typically designed to generate the types of nuclear deformations encountered by cells migrating through tight spaces, as would occur during immune or cancer cell intra- or extravasation. Missing from these systems is the ability to mimic the complex physical environment experienced by more quiescent, adherent cells such as epithelial or endothelial cells which, due to the topography and organization of the fibrous basement membrane on which they reside, are subjected *in vivo* to spatially non-uniform subcellular contact stresses exerted on their basal surfaces.

Systems that have targeted adherent cells to date have their own limitations. For instance, wavy surfaces have been used to demonstrate that cellular deformations at the peaks and valleys can be transmitted to the nuclei, resulting in changes in nuclear morphology, chromatin organization, and gene expression (Luciano *et al*, 2021; Pieuchot *et al*, 2018); however, the scale of the waviness in these systems leads to whole-cell rather than subcellular contact stresses. Subcellular, nuclear deformations have been reported in cells cultured on micropillars (Davidson et al, 2009, 2010; Badique et al, 2013; Ermis et al, 2016); however, these types of substrates fail to capture the anisotropy often present in the topography of basement membranes. In this context, we have been using microgroove substrates as idealized models of the anisotropic subcellular topography of the vascular basement membrane (Leclech *et al*, 2020). Our recent studies using these substrates have demonstrated their ability to control vascular endothelial cell morphology, cytoskeletal organization, and collective migration (Leclech *et al*, 2022, 2023).

In the present study, we extend the use of microgrooves to explore the impact of these topographic surfaces on cell nuclei. We show that microgrooves on the order of 5 µm in width, spacing, and depth elicit spontaneous, rapid, and robust nuclear deformations. Remarkably, these deformations include not only in-plane elongation but also in-depth penetration, ranging from nuclei that partially penetrate into the grooves to nuclei that are fully confined within the grooves. These 3D deformations are observed in a variety of cell types, and their extent can be controlled by modulating groove dimensions. We further show that these large-scale 3D nuclear deformations are dynamic with nuclei going into and out of the grooves cyclically and with no apparent DNA damage. Interestingly, AFM measurements demonstrate that these entry and exit cycles are accompanied by transient changes in perinuclear stiffness. While cell membrane protrusion into and adhesion to the groove surfaces are a prerequisite for nuclear penetration into the grooves, cytoskeleton-generated forces appear to surprisingly have a limited impact on this process. Finally, as a proof-of-concept of the potential relevance of these large-scale 3D nuclear deformations to health and disease, we show that myoblasts (muscle precursor cells) from patients with mutations in the *LMNA* gene cultured in the microgroove system exhibit abnormal nuclear deformations that can be detected and quantified using automated image analysis.

## Results

### 3D nuclear deformations in microgrooves are observed in various cell types

Microgroove substrates have been widely employed to generate cellular elongation and alignment, a phenomenon known as contact guidance (Leclech & Villard, 2020). We have recently studied the mechanisms underlying contact guidance in human umbilical vein endothelial cells (HUVECs) cultured on fibronectin-coated polydimethylsiloxane (PDMS) microgrooves (Fig. 1A) (Leclech *et al*, 2023). In the course of these investigations, we came across the observation that for a microgroove depth of ∼5 µm, the cell nuclei exhibited significant out-of-plane deformation and penetration into the grooves, in addition to the planar nuclear elongation and alignment associated with contact guidance (Fig. 1A). Importantly, unlike studies where nuclear elongation is driven by overall cell elongation on grooves or adhesive lines (McNamara *et al*, 2012; Bautista *et al*, 2019; Versaevel *et al*, 2012), the groove dimensions used here enabled confinement of the nucleus within the grooves while the rest of the cell remained free. These 3D nuclear deformations occurred very rapidly (as early as 30 min) after cell seeding and persisted for several days in culture, with the highest probability of occurrence between 2 and 24 h after cell plating. Additionally, nuclei of individual cells exhibited greater deformations than those of cell monolayers (data not shown). These initial observations prompted us to explore the use of microgrooves as a simple, robust platform to induce and analyze three-dimensional nuclear deformations in different adherent cells.

**Figure 1:**
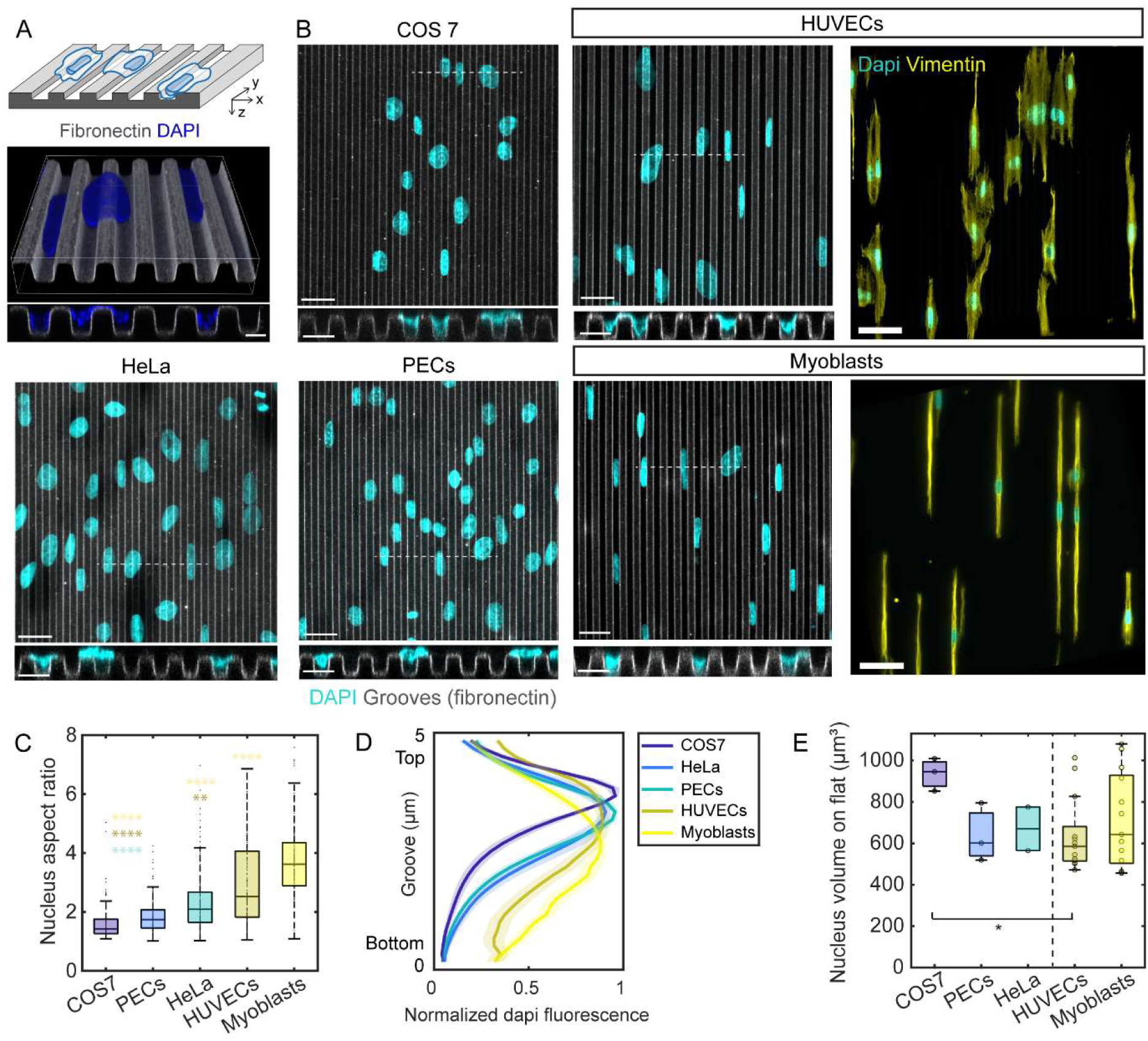
Nuclear deformations on microgrooves in different cell types. **(A)** Schematic of cell culture on microgroove substrates (top). Confocal microscopy 3D reconstruction and cross-section of nuclei (blue) on microgrooves visualized with fluorescent fibronectin (grey). Scale bar 5 µm (bottom). **(B)** Z-projection images (scale bar 25 µm) and cross-sections (scale bar 10 µm) from confocal stacks showing nuclei (stained with DAPI, blue) after 8 h of culture on microgrooves (grey, width = spacing = depth = 5 µm, denoted as 5×5×5 µm) for COS7 cells, HeLa cells, parietal epithelial cells (PECs), human umbilical vein endothelial cells (HUVECs), and myoblasts. Right panel shows immunostaining for vimentin to visualize cell shape. Scale bar 50 µm. **(C)** Quantification of the nucleus aspect ratio on microgrooves for the different cell types (n=109-231 cells from 3 independent experiments). **(D)** Quantification of normalized DAPI fluorescence intensity along groove depth. **(E)** Nuclear volumes of cells on control flat surfaces (n=2 to 13 independent experiments, 100 to 350 cells/condition). For all plots, one-way ANOVA, Dunn’s post-test (* p < 0.1; ** p < 0.01; **** p < 0.0001).

To test if these observations initially made on HUVECs were unique to endothelial cells, we cultured different cell types on microgroove substrates and analyzed the behavior of their nuclei (Fig. 1B). We observed different extents of nuclear elongation with nuclear aspect ratios ranging from 1.6 in COS 7 cells to 3.7 in myoblasts (Fig. 1C). Nuclear elongation was associated with at least partial nuclear penetration into the grooves for all cell types tested (Fig. 1B, cross-sectional views). To estimate the extent of nuclear penetration into the grooves, we quantified the average DAPI fluorescence intensity in different confocal z-planes in the direction of groove depth (Fig. 1D). The analysis revealed that HUVECs and myoblasts, which showed the highest nuclear in-plane elongation (Fig. 1C), also exhibited fluorescence intensity profiles that were shifted further towards the bottom of the grooves than those of the other cell types tested (Fig. 1D), indicative of deeper nuclear penetration into the grooves and hence more pronounced 3D deformations. Although this enhanced capacity for groove penetration did not correlate with nuclear volume (Fig. 1E), the most pronounced nuclear penetrations into the microgrooves were typically associated with highly elongated cells, which was particularly evident in the case of myoblasts (Fig. 1B, right panel). Because they exhibit the most pronounced 3D nuclear deformations, we opted to focus on HUVECs and myoblasts in all subsequent work.

### Different classes of nuclear deformations can be generated in microgrooves and modulated by groove dimensions

We have thus far seen that sufficiently deep microgrooves are capable of generating complex 3D nuclear deformations including both in-plane elongation associated with contact guidance and out-of-plane nuclear penetration into the grooves. For simplicity, we will henceforth refer to the combination of both of these types of deformation as “nuclear deformations”. Careful observation of confocal images of nuclei on microgrooves revealed the coexistence of different categories of deformations (Fig. 2A) associated with specific morphometric features (Fig. 2B). The “uncaged” category refers to nuclei suspended above the grooves that are elongated and oriented in the groove direction, remain flat with very little deformation in z, and exhibit morphological features close to nuclei on flat control PDMS surfaces (Fig. 2A,B). Note that this category was observed in HUVECs but not in myoblasts where all nuclei were at least partly deformed in z (Fig. S1A). On the other end of the deformation spectrum, nuclei can be fully confined within one groove (henceforth referred to as “caged” nuclei). These nuclei exhibit a very specific, highly elongated shape (aspect ratio of ∼5, Fig. 2B) associated with a significant decrease in projected area and loss of volume of ∼30% for HUVECs and ∼50% for myoblasts compared to nuclei on flat surfaces (Fig. 2B and Fig. S1B). The remaining nuclei in the population, termed “partly caged”, exhibit an intermediate level of deformation and thus lie partly within either one or two grooves and partly on a ridge (Fig. 2A). The partly caged nuclei are characterized by a highly tortuous contour as quantified by the solidity of the nuclear cross-section (Fig. 2A,B). Contrary to uncaged nuclei or nuclei on flat surfaces which are very flat and have smooth surfaces, caged and partly caged nuclei are thicker and exhibit surface wrinkles.

**Figure 2:**
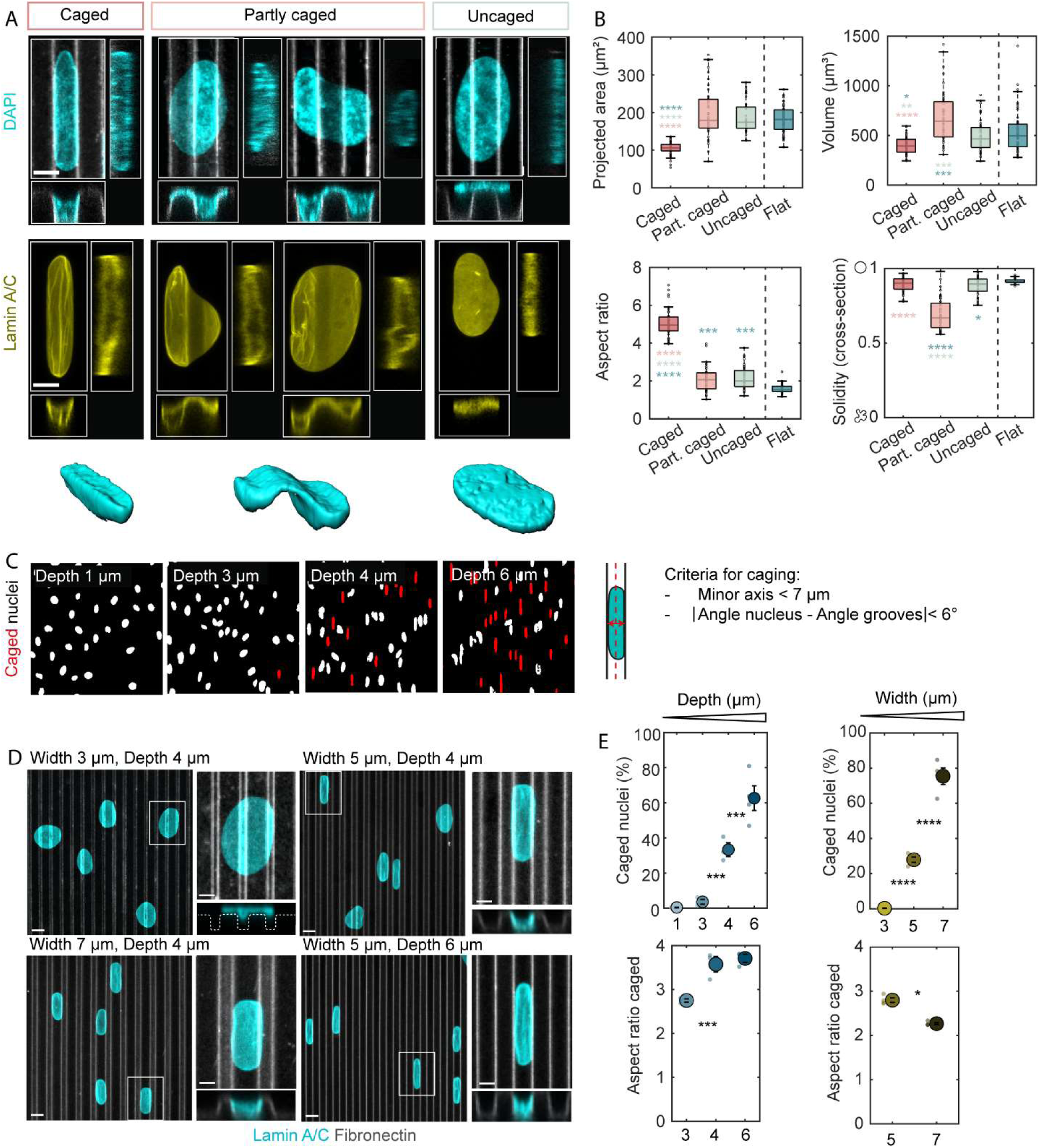
Characterization of nuclear deformations on microgrooves in HUVECs. **(A)** Z-projection images, cross-sections, and 3D reconstructions of the different classes of nuclei observed on microgrooves (5×5×5 µm), with DAPI (top) or lamin A/C staining (bottom). Scale bar 5 µm. **(B)** Morphological characterization of the different classes of nuclei observed on microgrooves and on control flat surfaces. Projected area and aspect ratio were quantified on z-projection images, solidity (tortuosity) on the cross-section images, and volume on 3D reconstructions. n=42-50 cells/category from 3 independent experiments. **(C)** Criteria for nuclear caging and output of the automatic detection of caged nuclei (red) for different groove depths (width = spacing = 5 µm). **(D)** Nuclei (stained with lamin A/C, cyan) on microgrooves (grey) of different dimensions. Scale bars 10 µm, 5 µm (zoom-ins). **(E)** Quantification of the percentage of caged nuclei and aspect ratio of caged nuclei for different groove depths (width = spacing = 5 µm) or groove widths (spacing = 5 µm, depth = 4 µm). Dots represent individual experiments and error bars represent standard errors of the mean (SEM). n=3 to 5 independent experiments. For all plots: one-way ANOVA, Fisher’s post-test (* p < 0.1; ** p < 0.01; *** p < 0.001, **** p < 0.0001).

One major advantage of the groove geometry relative to other systems within which nuclear deformations have been reported is the generation of the deterministic, reproducible, and rather simple nuclear shapes described above. Consequently, the specific morphometric features identified here for the different categories of deformations allow us to directly infer the 3D deformations from the 2D morphology of nuclei. In particular, caged nuclei can be automatically detected with a simple criterion on nuclear orientation and minor axis length (Fig. 2C). The percentage of caged nuclei, which can hence be easily obtained on conventional widefield images and with high-throughput, provides a simple and automatic quantification of the extent of nuclear deformation in microgrooves under different conditions.

We next investigated the dependence of nuclear deformations on microgroove dimensions. For groove width smaller than 4 µm, nuclei are unable to fully enter the grooves (one exception is myoblast nuclei, suggestive of higher deformability, Fig. S1C). Increasing groove width from 5 to 7 µm increases the percentage of caged nuclei at the expense of partly caged and uncaged nuclei but also decreases their elongation (Fig. 2D,E and Fig. S1C,D), suggesting that nuclei tightly conform to the groove space. For groove widths greater than 7 µm, the lateral confinement provided by the grooves is too low to impose significant deformation on the cell nuclei (data not shown). Similarly, groove depths below 3 µm are too small to elicit real out-of-plane penetration. Caging is first detected in 4 µm-deep grooves and increases with groove depth, coupled with increased nuclear elongation until saturation for 6 µm-deep grooves (Fig. 2D,E and Fig. S1C,D). In conclusion, in HUVECs and myoblasts the nuclear deformations described here occur over only a limited range of groove dimensions, namely 4-7 µm in width and spacing and 4-6 µm in depth. Within this range, the types and extent of nuclear deformations can be controlled by choosing the appropriate set of groove dimensions for the considered cell type.

We then proceeded to examine potential functional consequences of nuclear deformations in microgrooves. The EdU assay in HUVECs cultured on microgrooves of different depths revealed a slightly increased proliferation rate on microgrooves relative to cells on control flat surfaces (Fig. S2A). The observation of EdU-positive nuclei and the witnessing of dividing caged nuclei during live-cell recordings attest to the fact that nuclear deformations in microgrooves do not inhibit cell proliferation.

We next examined potential DNA damage by immunostaining against phosphorylated H2AX (pH2AX), a commonly used marker for DNA repair after double-strand breaks (Sharma *et al*, 2012). Contrary to the positive control (etoposide-treated cells), no nuclear pH2AX signal was visible in HUVECs on either flat substrates or microgrooves (Fig. S2B). In myoblasts, pH2AX-positive cells were occasionally observed; however, this appeared to be equally frequent on flat substrates as on microgrooves, suggesting that the DNA damage is not related to the physical constraints imposed by the grooves. In both cell types, no occurrence of nuclear envelope rupture could be observed (data not shown). Thus, microgrooves constitute a tunable, high-throughput platform to induce well-controlled yet non-detrimental nuclear deformations.

### Nuclear deformations on microgrooves are dynamic

To further characterize nuclear deformations in microgrooves, we sought to determine the dynamics involved in nuclear caging. To this end, we used Hoechst to visualize nuclei in live cells and recorded their dynamics over a period of 24 h. Elongated, caged nuclei could easily be identified in the recordings (Fig. 3A and Movies S1 and S2). Tracking individual nuclei revealed that caging is not permanent and that nuclei can go in and out of the grooves cyclically. Each nuclear trajectory (see colored arrowheads, Fig. 3A) can therefore contain caging phases separated by entry and exit phases (asterisks in Fig. 3A), during which nuclei exhibit the characteristic morphology of the “partly caged” category. In HUVECs, caging phases are often separated by uncaged phases (corresponding to either suspended or slightly deformed nuclei) while in myoblasts, the caging phases are predominant with nuclei quickly deforming while moving from one groove to another (Fig. 3A). More specifically, quantification shows that the caging phases are significantly longer in myoblasts (∼13 h) than in HUVECs (∼3 h) and that myoblasts correspondingly exhibit less frequent transitions among the different phases (Fig. 3B). Interestingly, HUVEC migration speed is significantly higher during the caging phase than during the other phases; this effect is not visible in the case of myoblasts since nuclei are caged for the great majority of the recording time (Fig. 3B). The observed differences in nuclear caging dynamics between HUVECs and myoblasts (short and cyclic in HUVECs vs. long and stable for myoblasts) are consistent with the different proportions of caged nuclei observed in fixed samples for these two cell types (cf: Figs. 2E and S1D).

**Figure 3:**
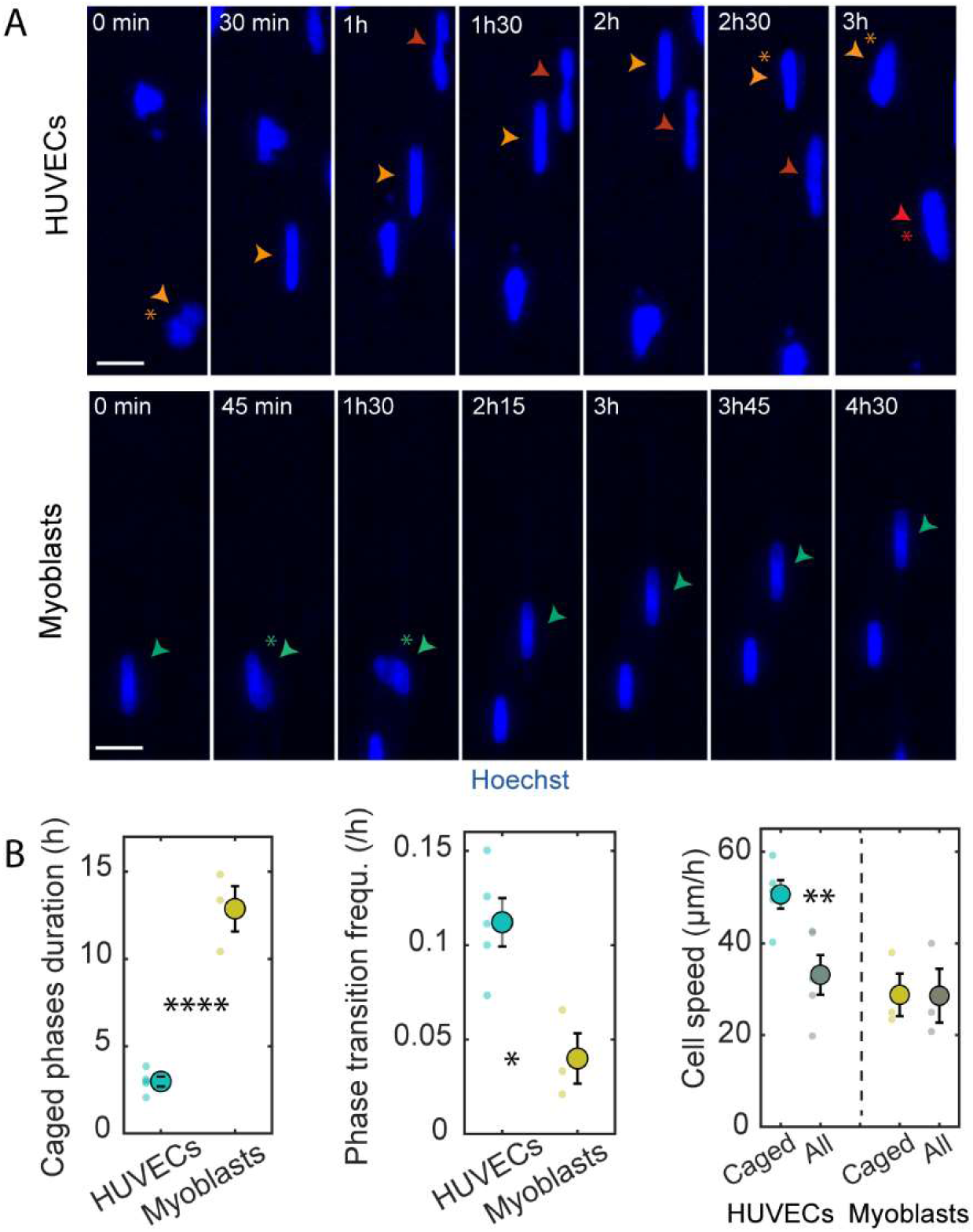
Dynamics of nuclear deformations on microgrooves. **(A)** Images extracted from time-lapse recordings of HUVEC or myoblast nuclei (stained with Hoechst, blue) on microgrooves (5×5×5 µm, vertical). Arrowheads follow the path of a single nucleus, and stars indicate uncaging phases. Scale bar 20 µm. **(B)** Quantification of the mean duration of caging phases, frequency of transitions between phases (caging-uncaging), and mean nucleus speed during caging phases compared to the mean speed of all nuclei at all times (“All”). Dots represent individual experiments, and error bars represent standard error of the mean (SEM). n=3 to 5 independent experiments. Student t test (2 groups) or one-way ANOVA, Fisher’s post-test (* p < 0.1; ** p < 0.01; **** p < 0.0001).

### Nuclear deformations in microgrooves are driven by cellular deformations

We next turned our attention to the mechanisms that underlie nuclear deformation in microgrooves. To this end, we started by examining the potential link between nuclear and overall cellular behavior. We first considered a large number of HUVECs and analyzed the relationship between the morphological parameters of a cell and its nucleus. Consistent with previous reports (Cantwell & Nurse, 2019), we observed a positive correlation between cell and nuclear areas (Fig. 4A, coefficient of correlation = 0.62) as well as cell and nuclear elongation (Fig. 4A, coefficient of correlation = 0.3). When focusing specifically on caged nuclei, it can be seen that while most caged nuclei are found in elongated cells, some caged nuclei are nevertheless observed in more round cells (group within the dashed outline in Fig. 4A). To better understand these observations, we performed time-lapse imaging of cells on microgrooves with both nuclear (Hoechst) and cytoplasmic membrane staining (CellMask dye) (Fig. 4B and Movies S3 and S4). These recordings revealed that nuclear movement overall follows cell movement, with nuclear exit from a groove preceded by the extension of a membrane protrusion into an adjacent groove (see arrows in Fig. 4B). In most cells, a significant change in cell shape appears to be simultaneously (at least within the 5 min interval between frames) accompanied by a change in nuclear shape (see “Cell1” in Fig. 4B, Movie S3). However, a time lag can also be observed between cell and nuclear movement, with nuclei remaining caged for a period of time even after the cell has moved significantly in another direction (see “Cell2” in Fig. 4B, Movie S4), explaining the “decorrelation” between cell and nuclear shape that is sometimes seen in fixed-cell images.

**Figure 4:**
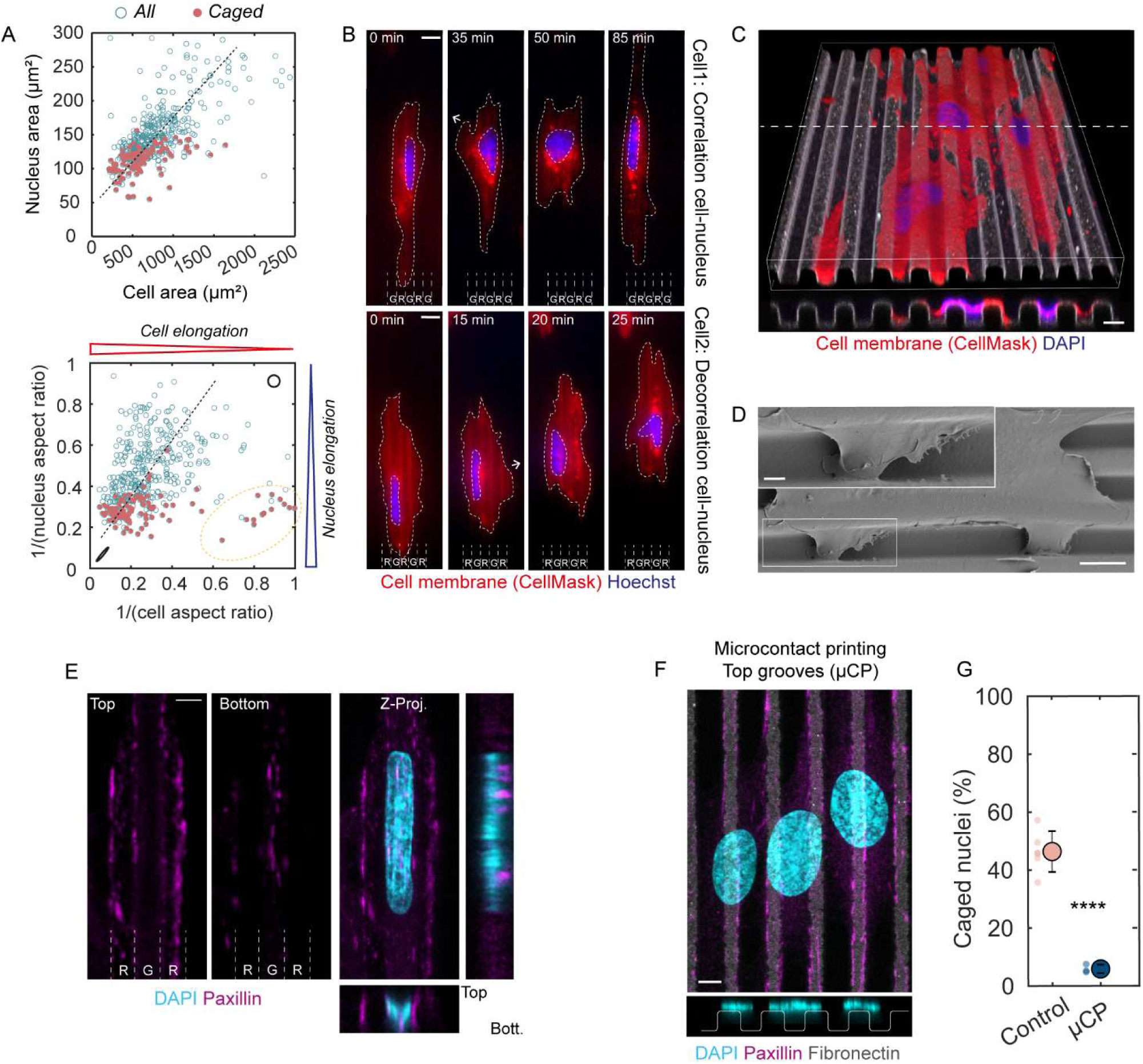
Influence of cell behavior on nuclear deformations on microgrooves. **(A)** Cell area vs. nuclear area and cell elongation vs. nuclear elongation. Dots represent individual cells (n=435 cells from 4 independent experiments), and caged nuclei are shown in magenta. Dotted lines are for the eye only and do not represent fits of the data. **(B)** Time-lapse images of two representative cells on microgrooves, with the cell membrane in red (CellMask) and nucleus in blue (Hoechst). The white arrows show the extension of a cell protrusion. Scale bar 10 µm. **(C)** 3D reconstruction and cross-section of HUVEC cell membranes (red, stained with CellMask) and nuclei (stained with Hoechst, blue) on microgrooves (5×5×5 µm). Scale bar 5 µm. **(D)** Electron microscopy of a cell on microgrooves. Scale bars, 10 µm, 2 µm (inset). **(E)** Immunostaining for paxillin (magenta) showing the presence of focal adhesions on both the ridge and groove surfaces. Scale bar 5 µm. **(F)** Immunostaining for paxillin (magenta), DAPI (blue), and fibronectin (grey) showing nuclear behavior on microgrooves where adhesion is restricted to the top of the ridges (microcontact printing). Scale bar 5 µm. **(G)** Quantification of the percentage of caged nuclei for an homogeneous fibronectin coating or microcontact printing experiments (µCP). Dots represent individual experiments and error bars represent standard deviations. n=3 to 6 independent experiments. One-way ANOVA, Fisher’s post-test (**** p < 0.0001).

In light of the decisive role that the cell seemingly has on nuclear behavior, we then proceeded to examine the cell membrane on microgrooves more carefully. In 3D reconstructions of either confocal or electron microscopy images, we observed large deformations of the cell membrane along the groove walls down to the bottom of the grooves (Fig. 4C,D). Staining for focal adhesions (paxillin) confirmed that cells establish adhesions both on the ridge surfaces and at the bottom of the grooves, below caged nuclei (Fig. 4E). To test if cell membrane protrusion into the grooves and the formation of focal adhesions within the grooves was required for nuclear deformations, we used microcontact printing to restrict the fibronectin coating to the ridges while the remaining groove surfaces remained uncoated (and thus non-adhesive) (Fig. 4F). In this case, nuclear caging was essentially abolished (Fig. 4G), indicating that cell membrane protrusion into and adhesion to the bottom of the grooves is necessary for subsequent nuclear entry into the grooves.

### An intact cytoskeleton is not necessary for nuclear caging in microgrooves

Because it links focal adhesions present at the bottom of the grooves to the nucleus, we hypothesized that the cell cytoskeleton may provide pulling forces that enable nuclear entry into the grooves. We first analyzed the 3D organization of the three main cytoskeletal networks: actin filaments, intermediate filaments (vimentin), and microtubules around caged nuclei in HUVECs (Fig. 5A). To this end, we quantified the localization of each cytoskeletal network relative to the nucleus by plotting its average fluorescence intensity as a function of groove depth (z) (Fig. 5B). While some actin filaments were present at the ridge level, the most prominent actin stress fibers localized below caged nuclei, indenting the nuclear envelope in some cases. In contrast, a dense vimentin network was visible above the nucleus but was largely absent from the bottom of the grooves. Microtubules exhibited an intermediate localization, mainly present above the nucleus but also visible on the side and occasionally below caged nuclei (Fig. 5A,B). This cytoskeletal organization was specific to caged nuclei, as illustrated by the different intensity profiles observed for caged and uncaged nuclei in Fig. 4B. Similar results were found for myoblast caged nuclei (Fig. S3A).

**Figure 5:**
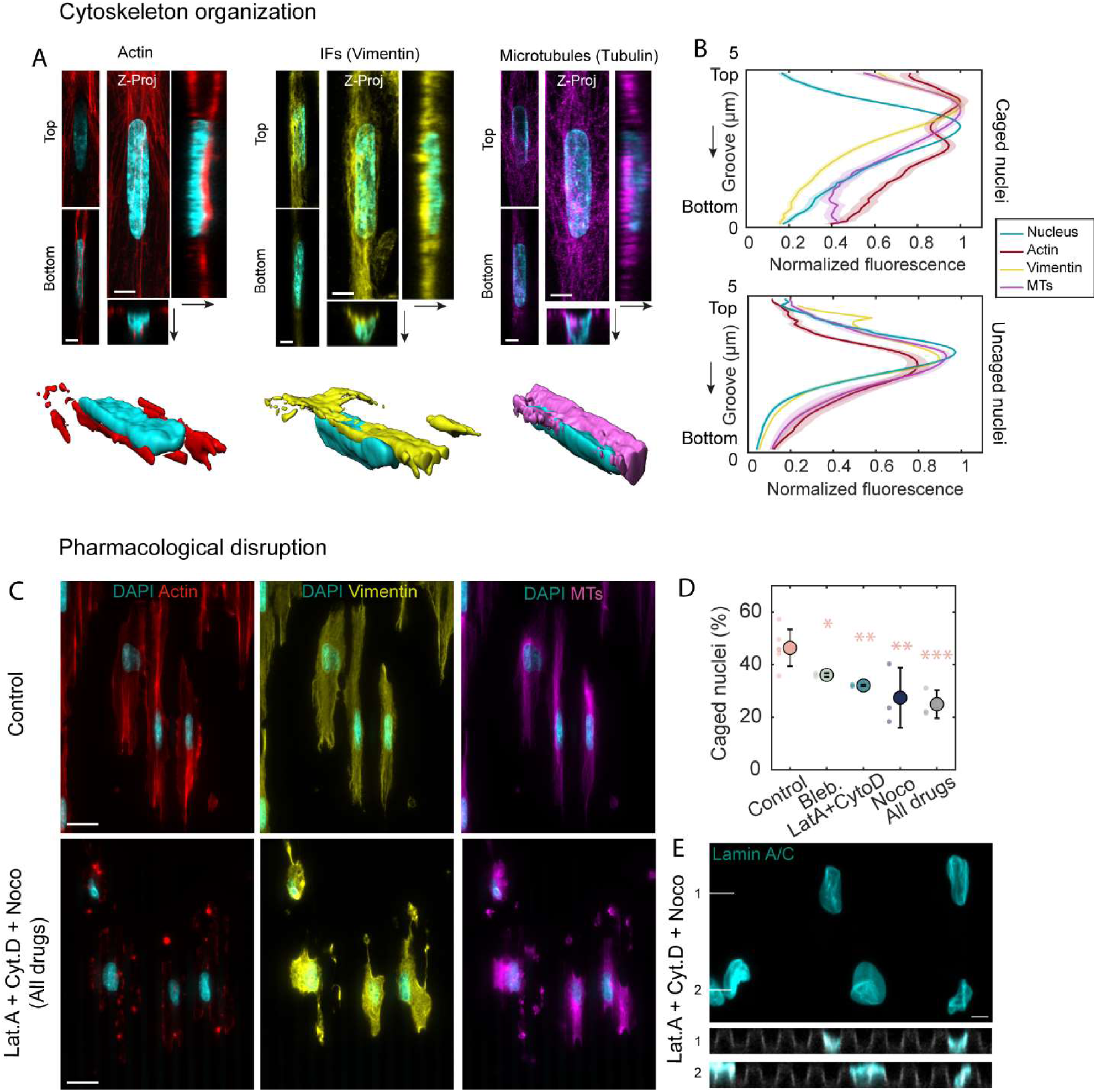
Organization and role of the cytoskeleton in nuclear deformations on microgrooves. **(A)** Images from confocal stacks and 3D reconstructions showing the organization of actin (red), intermediate filaments (vimentin, yellow), and microtubules (magenta) around caged nuclei. Scale bars 5 µm. **(B)** Quantification of the normalized fluorescence intensity for the three cytoskeletal networks and the nucleus as a function of depth (from top to bottom of the grooves) for caged (top) and uncaged (bottom) nuclei. **(C)** Immunostaining for actin (red), intermediate filaments (vimentin, yellow), and microtubules (magenta) in control cells or cells treated with latrunculin A (Lat.A), cytochalasin D (Cyto.D), and nocodazole (Noco) on microgrooves (5×5×5 µm, vertical). Scale bar 20 µm. **(D)** Quantification of the percentage of caged nuclei for different pharmacological treatments: control (DMSO), blebbistatin (Bleb.), latrunculin A and cytochalasin D (Lat.A + Cyto.D), nocodazole (Noco), or latrunculin A + cytochalasin D + nocodazole (All drugs). Dots represent individual experiments and error bars represent standard deviations. n=3 to 6 independent experiments. One-way ANOVA, Fisher’s post-test (* p < 0.1; ** p < 0.01; *** p < 0.001). **(E)** Z-projection and cross-sections of nuclei stained for lamin A/C and treated with latrunculin A + cytochalasin D + nocodazole. Scale bar 5 µm.

To test the involvement of the different cytoskeletal networks in nuclear caging, HUVECs were incubated with different pharmacological agents: blebbistatin (100 µM) to decrease cell contractility, latrunculin A (10 nM) plus cytochalasin D (20 nM) to disrupt both cortical actin and actin stress fibers, or nocodazole (200 nM) to disrupt microtubules. The cytoskeleton-disrupting agents were administered directly upon cell seeding in order to avoid confusing effects on nuclear entry or nuclear exit from the grooves. Surprisingly, none of these treatments completely abolished nuclear penetration into the grooves. Microtubule disruption had the largest effect, but even then, a minimum of ∼30% of the nuclei remained caged (Fig. 5D). To avoid to the extent possible a potential compensatory effect of any one of the cytoskeletal networks on the others and to target a minimalist system composed only of the cell nucleus, membrane, and cytoplasm, we incubated the cells with a cocktail of latrunculin A, cytochalasin D, and nocodazole (20 nM, 40 nM and 400 nM, respectively) (Fig. 5C). While the effect of this treatment was drastic on the cells with complete loss of contact guidance (Fig. 5C) as well as on nuclear morphology with more irregular shapes and wrinkling of the nuclear envelope, nuclear penetration into the grooves was not abolished (Fig. 5D, cross-sections). Similar results were obtained with myoblasts (Fig. S3B). These findings suggest that nuclear caging can occur in the absence of an intact cytoskeleton and that the cytoskeleton plays an accessory role in the process.

We further tested the involvement of the cytoskeleton by using myoblasts with a mutation in the KASH domain of Nesprin-1, a member of the Linker of Nucleoskeleton and Cytoskeleton (LINC) complex which links the actin cytoskeleton to the nuclear envelope (Fig. S3C). The mutation did not modify the occurrence of caging, suggesting that forces transmitted by the LINC complex are indeed not essential for nuclear caging.

To gain further insight into the minimal elements needed to obtain nuclear caging, we seeded isolated nuclei from myoblasts on microgrooves and observed their behavior after a couple of hours in culture. Interestingly, although full caging was never observed during that time, partial penetration of the nuclei inside the grooves was visible (Fig. S3D), suggesting that a part of the nuclear deformations on microgrooves originates from a combination of interaction with the substrate and intrinsic mechanical properties of the nucleus.

In summary, the results described above indicate that cytoskeleton-mediated pulling or pushing forces are not essential for nuclear caging, highlighting the robustness of this process. Rather, cell membrane adhesion and subsequent protrusion into and spreading on the microgrooves appear to provide a sufficient driving force for nuclear deformation. The cell cytoskeleton may subsequently actively reinforce this process.

### Nuclear deformations in microgrooves are associated with changes in perinuclear stiffness

For a given applied stress, nuclear deformation is determined by the mechanical properties of the nucleus. To probe the stiffness of nuclei in different states of deformation on microgrooves, we performed AFM experiments on both HUVECs and myoblasts with Hoechst-stained nuclei to identify both the position of the nucleus and the category of nuclear deformation (Fig. 6A). In these experiments, one measurement was made every two pixels over the entire nuclear area. The Young’s modulus was then extracted from the different force-distance curves in accordance with the Hertz model (see Methods for details) and averaged over the perinuclear area. While a possible contribution of the cell membrane and the cytoskeleton present above the nucleus cannot be completely excluded, we believe that this protocol allows us to obtain an acceptable approximation of the nucleus stiffness itself because: i) the nucleus is very closely apposed to the cell membrane in these very flat cells (see Fig. 4C), and ii) the measurements were made with a force setpoint that provides an indentation of 0.5 to 1 µm which corresponds to a significant portion of the cell height, thereby ensuring some probing of the nucleus below the membrane. In light of these arguments, the measured stiffness of the nuclear region (membrane, cortex, and nucleus) will henceforth be simply referred to as “perinuclear stiffness”.

**Figure 6:**
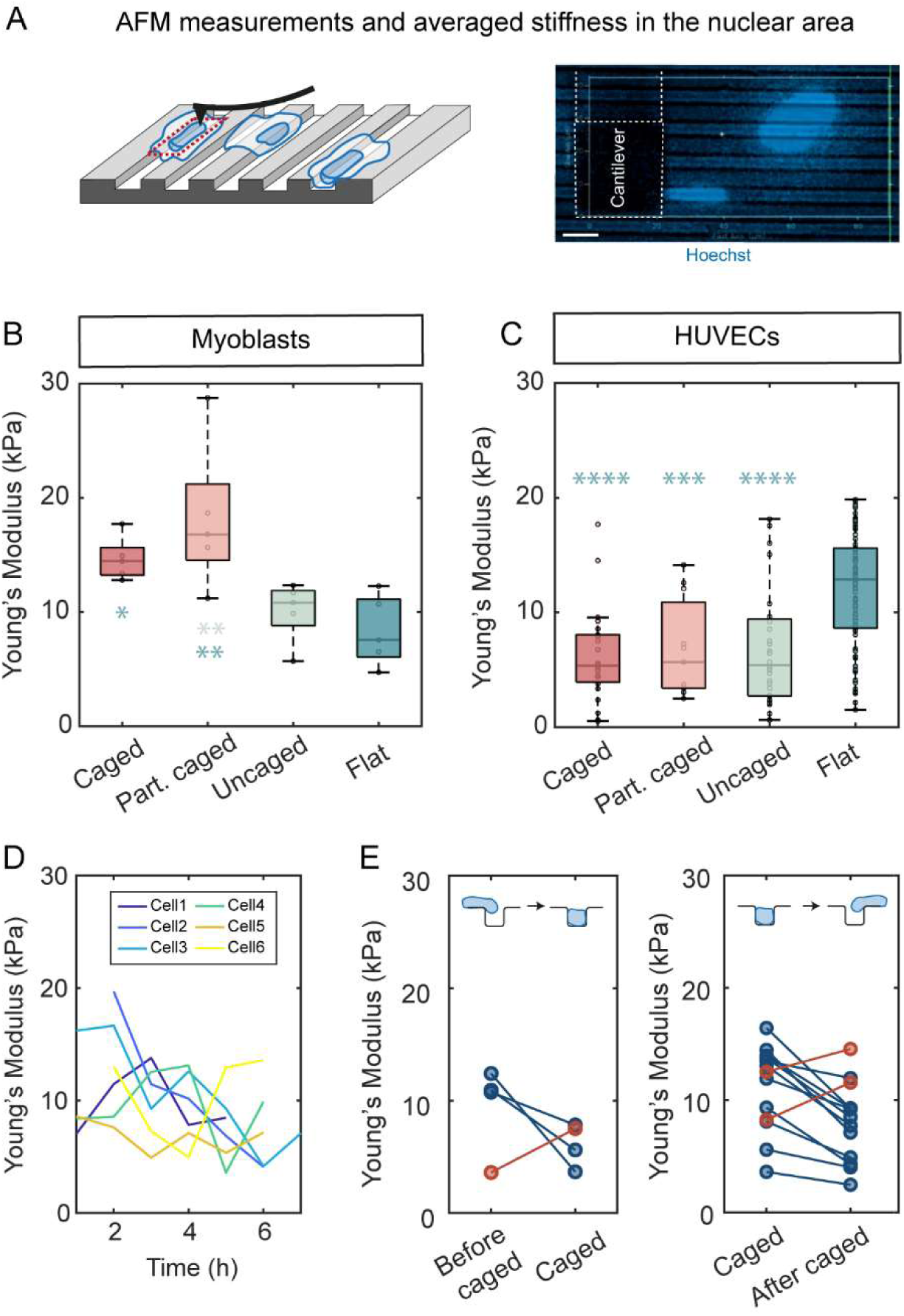
Nuclear deformations on microgrooves are associated with changes in nuclear stiffness. **(A)** Schematic of the atomic force microscopy (AFM) measurements, and image from the experiments showing the AFM cantilever from top and the nuclei stained with Hoechst. Scale bar 10 µm. **(B,C)** Young’s modulus of nuclei in different configurations of deformations on microgrooves or on flat PDMS surfaces, in myoblasts (top) or HUVECs (bottom). One-way ANOVA, Fisher’s post-test (* p < 0.1; ** p < 0.01; *** p < 0.001; **** p < 0.0001). **(D)** Evolution of the Young’s Modulus of 6 different HUVECs nuclei with time. **(E)** Evolution of the Young’s Modulus of different HUVECs nuclei during transitions from uncaged to caged (left) or caged to uncaged (right). The red dots and lines correspond to nuclei that exhibit behavior different from the main trend.

We first measured nuclei in different categories of deformation (caged, partly caged, or uncaged) as well as nuclei on flat surfaces. In myoblasts, caged and partly caged nuclei were significantly stiffer (Young’s moduli of 14.6 ± 1.9k Pa and 18.2 ± 6.4 kPa, respectively) than nuclei on flat surfaces (8.3 ± 3 kPa) (Fig. 6B). This result may initially appear counter-intuitive given the high deformability of this cell type’s nuclei; however, it may reflect nuclear structural remodeling and compaction as a result of caging (cf: Fig. S1B). In contrast, HUVEC nuclei on microgrooves were softer overall than those on flat substrates but did not exhibit significant stiffness differences among the three deformation categories on microgrooves, principally due to the large spread in the measurements (Fig. 6C). We hypothesized that this variability may be linked to the highly dynamic cell and nuclear behavior in this cell type. To test this hypothesis, we made successive AFM measurements of single nuclei over time (every hour during 4 to 8 h) and found that nuclear stiffness varies significantly in time within one cell (Fig. 6D). Fortunately, we were able to also monitor caging transitions during the course of these measurements, with nuclei going into or out of the grooves. Interestingly, both of these transitions were associated with rapid changes in nuclear stiffness, most commonly transient nuclear softening (Fig. 6E). These results suggest the existence of a cycle of changes in nuclear stiffness associated with the cycle of nuclear deformations. However, whether these changes in mechanical properties drive the observed nuclear deformations or are a consequence of these deformations remain to be determined.

### Myoblasts with *LMNA* mutations exhibit abnormal nuclear deformations on microgrooves

Various pathologies such as laminopathies and certain types of cancer are associated with alterations in nuclear mechanical properties (Zwerger *et al*, 2011). In light of the observed changes in nuclear stiffness that accompany nuclear deformations in microgrooves, we hypothesized that the microgroove platform may be useful in unveiling abnormalities in nuclear mechanical properties. To test this hypothesis, we cultured normal myoblasts as well as myoblasts derived from patients with muscular laminopathies on microgrooves. More specifically, we studied 3 different heterozygous mutations in the *LMNA* gene (which codes for A-type lamins) responsible for severe congenital muscular dystrophies. All mutant cells exhibited drastically different nuclear deformations on microgrooves compared to WT cells (Fig. 7A). Overall, the incidence of nuclear caging was smaller in mutants which instead exhibited more partly caged, highly tortuous nuclei (Fig. 7B,D). When caging occurred, the nuclei of mutant myoblasts were significantly more elongated those of the WT cells (Fig. 7C). Time-lapse imaging showed that mutant cells tended to migrate faster than WT cells and to exhibit faster, shorter but more frequent caged phases (Fig. S4). Interestingly, the microgroove system allowed the detection of differences among mutations. For instance, the ΔK32 mutation was associated with very long caged nuclei, while cells with the L380S mutation exhibited very tortuous partly caged nuclei. Overall, nuclei from patient cells appear to be more deformable than WT cells and are seemingly unable to maintain their structural integrity in response to the physical constraints imposed by the microgrooves. Because they mechanically challenge nuclei, microgrooves are able to reveal functional abnormalities that can be automatically detected and quantified, paving the way for potential diagnostic applications.

**Figure 7:**
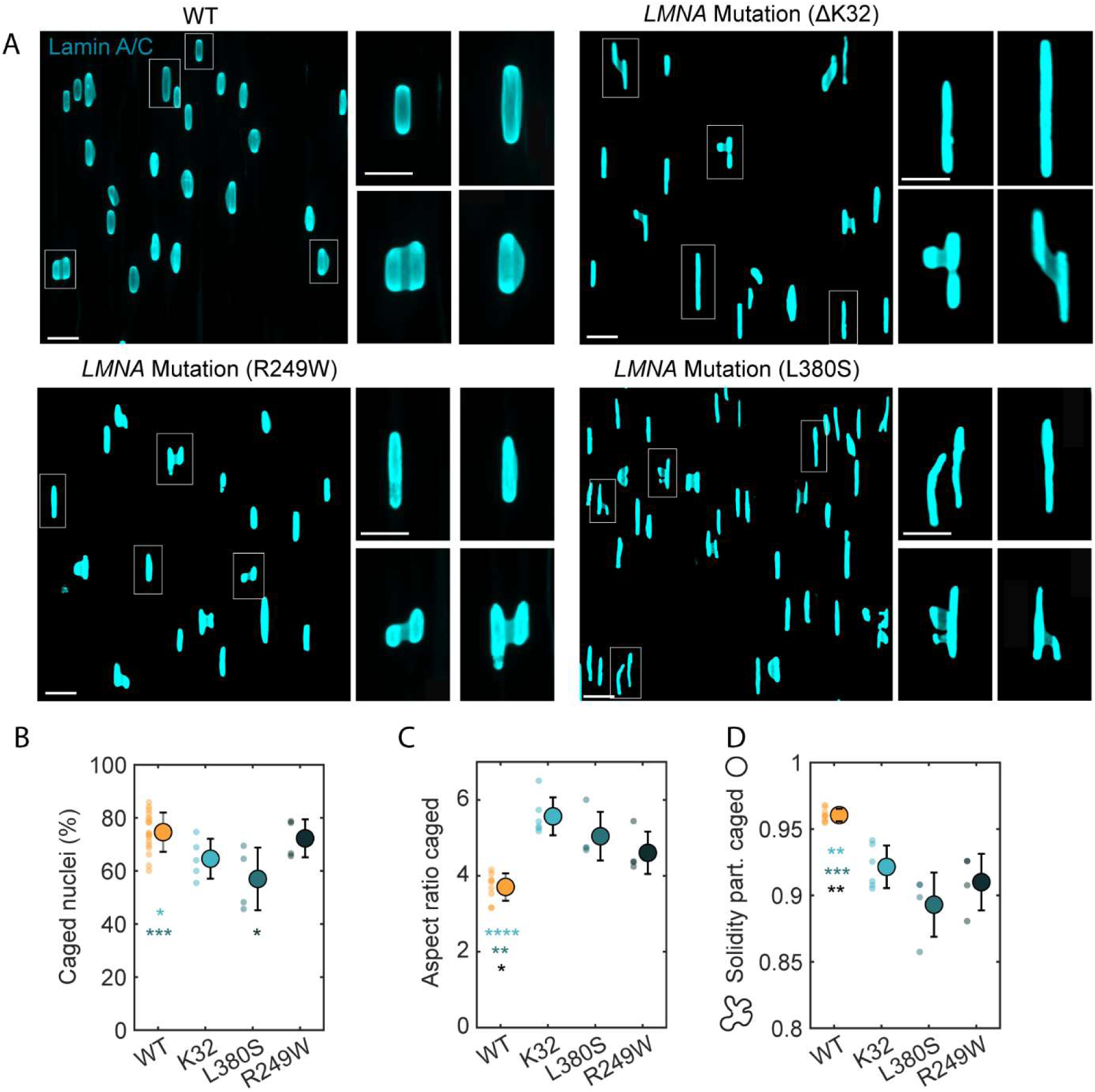
Myoblasts with LMNA mutations exhibit abnormal nuclear deformations on microgrooves. **(A)** Nuclei from WT myoblasts or myoblasts carrying mutations in the LMNA gene (ΔK32, R249W or L380S) on microgrooves, immunostained for lamin A/C. Insets show zoom-ins on representative nuclei. Scale bars 30 µm, 20 µm (insets). **(B)** Quantification of the percentage of caged nuclei. **(C)** Aspect ratio of caged nuclei. **(D)** Solidity of partly caged nuclei. Dots represent individual experiments and error bars represent standard deviations. n=4 to 9 independent experiments. One-way ANOVA, Fisher’s post-test (* p < 0.1; ** p < 0.01; *** p < 0.001; **** p < 0.001).

## Discussion

In this study, we used microgroove substrates to investigate the nuclear deformations that adherent cells might experience in response to micrometric anisotropic topographical cues stemming from their underlying basement membrane. For instance, in the vascular system, endothelial cells are subjected to topographical constraints at different scales, including micrometric undulations of the vessel wall. Nuclear deformations observed here *in vitro* could therefore occur *in vivo*, where it might, in addition to apical shear forces, influence endothelial cells intranuclear organization, transcriptomic activity or activation of downstream signaling events.

The principal novelty of the current work is that in addition to the nuclear elongation in the direction of the microgrooves associated with cellular contact guidance that has been described in several previous studies, we also report significant out-of-plane deformations, with partial to full “caging” of nuclei inside the grooves. These 3D nuclear deformations occur for a specific range of groove dimensions (width, spacing and depth between 4 and 7 µm), which allow for a specific, physical confinement of the nucleus while the remainder of the cell remains free. Different categories of nuclear deformation can be generated using the microgroove system, and the proportion of the different categories can be precisely controlled by varying the groove dimensions, rendering the system easily adaptable to different experimental settings and cell types. In contrast to micropillar substrates which induce complex and random nuclear deformations, the principal advantage of the groove geometry lies in its ability to generate these deterministic and reproducible nuclear shapes, such as caged nuclei. This categorization serves as a simple reference metric for quantifying nuclear deformations under different conditions.

In adherent cells, cytoskeletal forces are usually the primary drivers of nuclear deformation (Tusamda Wakhloo et al, 2020; Versaevel et al, 2012). The specific organization of cytoskeletal elements around caged nuclei with actin stress fibers principally below the nuclei and intermediate filaments and microtubules above and at the level of the nuclei initially suggested that actin cables might serve to pull the nuclei into the grooves from below while intermediate filaments and microtubules may push down from above. Surprisingly, however, extensive pharmacological disruption of the actin and microtubule cytoskeleton failed to abolish nuclear entry into and caging within the grooves. It should be noted that although intermediate filaments were not targeted directly in the pharmacological disruption studies, vimentin staining revealed that intermediate filaments were nevertheless strongly disrupted by the relatively high doses of actin- and microtubule-disrupting drugs used. We note also that intermediate filaments have been shown in other studies to protect and maintain nuclear shape rather than induce nuclear deformation (Neelam et al, 2015; Patteson et al, 2019). In another albeit different experimental setting, it has been reported that upon microdissection from the cell body, nuclei retain their original shape, indicating that nuclear deformation can be decoupled from instantaneous cell shape-dependent cytoskeletal forces (Tocco et al, 2018). Taken together, the various observations described above have led us to conclude that nuclear caging in our system only weakly relies on cytoskeleton-generated forces. On the other hand, we showed that cell membrane deformation and adhesion into the groove is necessary for subsequent nuclear caging. We therefore propose a model in which cellular adhesion forces generate sufficiently large downward-acting forces on the cell membrane to push the nucleus into the groove and induce nuclear caging.

A particularly intriguing attribute of the nuclear confinement reported here is its dynamic nature whereby nuclei repeatedly and cyclically enter and exit the microgrooves. Such dynamics have not been reported in other experimental settings that induce nuclear deformations such as micropillar arrays and thus appear to be another distinguishing feature of the microgroove system. Interestingly, the observed dynamics vary among cell types, with HUVECs exhibiting shorter but more frequent caging phases compared to myoblasts which remain caged for significantly longer periods of time (13 h vs. 3 h on average). Consistent with this observation, in fixed-cell samples, ∼80% of myoblast nuclei on 5 µm-deep grooves are caged vs. ∼50% for HUVECs. These differences in nuclear deformation dynamics may stem from intrinsic differences in nuclear mechanical properties between the two cell types, as suggested by the AFM data showing that myoblast nuclei on flat surfaces are softer (Young’s modulus of ∼8 kPa) than those of HUVECs (∼12 kPa).

In myoblasts, the prolonged caging appears to be accompanied by nuclear stiffening. Although we did not observe a significant change in DAPI intensity in caged nuclei, this may reflect a level of nuclear compaction. In HUVECs, the dynamic caging and uncaging events are associated with transient changes in nuclear mechanical properties that most commonly take the form of perinuclear softening. The structural basis for these changes remains unknown but may be linked to dynamic reorganization and modification of the lamin network in the nuclear envelope. Whether this form of mechanical adaptation is a cause or a consequence of nuclear deformation remains to be established, i.e. while these nuclear stiffness changes might passively arise from structural modifications associated with nuclear deformations, they may also more actively enhance the deformability of nuclei to facilitate their entry into or exit from the grooves, akin to cancer cell nuclei softening during transendothelial migration (Roberts *et al*, 2021). Most cell types, including the muscle and vascular cells studied here, are constantly mechanically challenged by their extracellular environment. Nuclear mechanical adjustments on short time scales and the underlying mechanisms are therefore of great interest but remain very poorly understood, principally due to a lack of adapted tools. In this context, the combination of microgrooves with dynamic AFM measurements as conducted here provides a powerful approach for studying the process of dynamic cell and nuclear deformation.

Despite the large-scale nuclear deformations and loss of nuclear volume reported here, microgroove-induced nuclear caging does not elicit visible DNA damage, at least in cells without nuclear envelope mutations. This may be attributable to the dynamic nature of nuclear caging which shields the nucleus from prolonged mechanical stress. The mechanical adaptation of the nuclei (softening) upon entering or exiting the grooves may also help. It is also possible that the cell types tested express relatively high levels of lamins, allowing the nuclear envelope to resist such deformations without rupturing. Overall, this result contrasts with other systems such as compression devices or microfluidic channels with constrictions which often generate nuclear envelope rupture and DNA damage (Nader et al, 2021; Raab et al, 2016). In such systems, it is important to note that the mechanical stresses are actively imposed onto the nuclei whereas the deformations in the case of the microgrooves are “self-imposed”. This second, more physiological setting could allow the cells to adapt efficiently (i.e. with less damage) to the physical and mechanical constraints present in the extracellular environment.

An association between pathologies and cellular mechanics has long been proposed and is now increasingly supported by evidence linking changes in cellular or nuclear mechanics to disease development (Zwerger *et al*, 2011; Harris *et al*, 2019). Mechanical biomarkers therefore constitute a promising approach for the detection of some of these pathologies, and efforts have been made to develop experimental systems exploiting these novel biomarkers (Mao & Huang, 2012). Various types of microfluidic devices based principally on deformability cytometry (constriction deformability or fluid shear deformability (Chen et al, 2022)) have been used to assess cell mechanics. In this context, because of its simplicity, robustness, and high-throughput, microgroove substrates can constitute a valuable tool to functionally test cell deformability and more specifically that of their nuclei. As a proof of concept of this idea, we were able to use the microgrooves to unveil significant variations in nuclear shapes between healthy myoblasts and myoblasts carrying three different mutations in the *LMNA* gene that cause severe muscular dystrophies (laminopathies). Thus, microgrooves constitute a promising tool to functionally probe nuclear integrity in laminopathies, where determining the pathogenicity of diverse mutations can be challenging. Further investigations targeting other pathologies, such as cancer, will test the applicability range of this system, potentially opening new avenues for diagnostic and therapeutic applications.

In conclusion, we propose the novel use of microgrooves as a simple, tunable, and high-throughput platform to induce and study nuclear deformations and mechanics in various cell types. As previously detailed, this system offers several advantages and unique characteristics that can nicely complement other existing systems in this field. In addition, we believe it has the potential to serve as a functional diagnostic platform for the detection of pathological alterations in nuclear structure and mechanics.

## Methods

### Fabrication of microgrooved substrates

The original microstructured silicon wafer was fabricated using photolithography by direct exposure of a layer of SU8 photoresist (MicroChem, USA) with a µPG machine (Heidelberg Instruments). After exposure to trichloro(1H,1H,2H,2H-perfluorooctyl)silane (Sigma) vapor for 1 h, the original wafer was used to create polydimethylsiloxane (PDMS Sylgard 184, Sigma Aldrich, ratio 1:10) replicates. To create the final coverslip on which the cells were cultured, liquid PDMS was spin coated at 1500 rpm for 30 s on the PDMS mold. Before reticulation overnight at 70°C, a glass coverslip was placed on top of the PDMS layer. After reticulation, the glass coverslip attached to the microstructured PDMS layer was gently demolded with a scalpel and isopropanol to facilitate detachment. Microstructured coverslips were then sonicated for 10 min in ethanol for cleaning and finally rinsed with water.

### Cell culture

Microgroove substrates were incubated for 1 h with 50 μg/ml fibronectin solution (Sigma F1141) at room temperature after a 30 s plasma treatment. Mixing of fibronectin with fluorescent fibrinogen 647 (Thermofisher F35200) was used to visualize fibronectin localization. All cell types were cultured at 37°C in a humidified atmosphere of 95% air and 5% CO_2_. Before culture, cells were detached with trypsin (Gibco, Thermo Fisher Scientific) and seeded onto microgroove coverslips at densities of 30,000-50,000 cells/cm^2^ for 2 to 24 hours.

*Endothelial cells:* Human umbilical vein endothelial cells (HUVECs, Lonza) in passages 4-8 were cultured in EGM2-MV medium (Lonza).

*Myoblasts*: Muscle biopsies were obtained from the Bank of Tissues for Research (Myobank, a partner in the EU network EuroBioBank, Paris, France) in accordance with European recommendations and French legislation. Following muscle biopsies, muscle cell precursors were immortalized as previously described (Mamchaoui *et al*, 2011). Control myoblasts were immortalized from one healthy subject. Immortalized human myoblasts carrying the following heterozygous mutations responsible for severe congenital disorders were also used: *LMNA* c.94_96delAAG, p.Lys32del (hereafter referred to as ΔK32), *LMNA* p.Arg249Trp (hereafter referred to as R249W), *LMNA* p.Leu380Ser (hereafter referred to as L380S) and *SYNE-1* homozygous c.23560 G<T, p.E7854X leading to a stop codon in exon 133 and deletion of the carboxy-terminal KASH domain (hereafter referred to as Nespr-1ΔKASH). Myoblasts were cultured in growth medium consisting of 1 vol 199 Medium/4 vol DMEM (Life Technologies, Carlsbad, CA, USA) supplemented with 20% fetal calf serum (Life technologies, Carlsbad, CA, USA), 5 ng/mL hEGF (Life Technologies, Carlsbad, CA, USA), 0.5 ng/mL βFGF, 0.1 mg/mL dexamethasone (Sigma-Aldrich, St. Louis, MO, USA), 50 μg/mL fetuin (Life Technologies, Carlsbad,CA, USA), 5 μg/mL insulin (Life Technologies, Carlsbad, CA, USA), and 50 mg/mL Gentamycin (Gibco™, Life Technologies, Carlsbad, CA, USA).

*Parietal Epithelial Cells*: Primary mouse Parietal Epithelial Cells (mPECs) (Kabgani *et al*, 2012) (Material Trade Agreement from M. Moeller, University Hospital of the Aachen University of Technology) were isolated using transgenic mouse lines and fluorescence-activated cell sorting (FACS). mPECs were cultured in supplemented ECBM (supplement kit Promocell) with Pen-Strep and 20% FBS until 70% of confluence before passaging.

*HeLa and COS-7 cells* were purchased from ATCC and cultured in DMEM-glutamax with 10% FBS and Pen-Strep.

### Cytoskeletal disruption

The following cytoskeletal pharmacological agents were used: blebbistatin (Sigma B0560) at 100 µM, latrunculin A (Millipore 428026) at 10 nM (HUVECs) or 1 µM (myoblasts), cytochalasin D (Sigma C2618) at 20 nM, (HUVECs) or 1 µM (myoblasts) and nocodazole (Sigma M1404) at 0.2 µM. When used together, the following concentrations were used: latrunculin A 20 nM, cytochalasin D 40 nM and nocodazole 0.4 µM. Drugs were incubated directly with the cells upon seeding and left for 90-120 min. Controls were treated with the equivalent DMSO concentration.

### EdU assay for cell proliferation

Cell proliferation was assessed using the “Click-iT Plus EdU imaging” kit (ThermoFisher C10640) after 24 hours of culture, with an EdU incubation time of 7 h.

### Microcontact printing

Flat PDMS blocks were used as stamps. After incubation with fibronectin (50 μg/ml) mixed with fibrinogen 488 in PBS for 1 h at room temperature, stamps were rinsed once with water and dried with an air gun. Microgroove coverslips were plasma-activated and were rapidly placed in contact with the stamps for 3 min under a 30 g mass, which deposited fibronectin only on the groove ridges. Cells were then cultured on the surfaces of the microgrooves as described above.

### Isolation of cell nuclei

All steps were carried out at 4 °C. After PBS wash, nuclei were isolated in a homogenization buffer containing 10 mM HEPES and 1mM DTT. Cells were homogenized with approximately 25 strokes using a Dounce homogenizer, collected in homogenization buffer and loaded over a 30% sucrose gradient. The samples were mixed thoroughly by inverting and then centrifuged at 800g for 10 min, yielding a crude nuclear pellet. The pellets were resuspended in 10 mM HEPES/1 mM DTT, centrifuged again and resuspended in HEPES/DTT. Nuclei were then plated on microgrooves and cultured for 3 to 6 h.

### Immunostaining

Culture coverslips were fixed with 4% paraformaldehyde (Thermo Fisher) in PBS for 15 min. After 1 h in a blocking solution containing 0.25% Triton and 2% bovine serum albumin (BSA), cultures were incubated for 1 h at room temperature with primary antibodies as follows: mouse anti-laminA/C (Sigma SAB4200236), mouse anti-phospho-histone H2AX (Merck 05-636), mouse anti-paxillin (MA5-13356, Thermofisher), mouse anti-vimentin (ab8069, Abcam) or rabbit anti-vimentin (ab92547, Abcam), mouse anti α-tubulin (Sigma, T5168), All antibodies were diluted 1/400 - 1/200 in a solution containing 0.25% Triton and 1% BSA. Coverslips were washed three times with PBS and incubated for 1 h at room temperature with Alexa Fluor 555-conjugated donkey anti-rabbit antibody (ab150074, Abcam) or Alexa Fluor 488-conjugated donkey anti-mouse antibody (ab150105, Abcam) and DAPI. When needed, actin was stained during this last step using phalloidin (LifeTechnologies).

### Microscopy

#### Epifluorescence and confocal microscopy of fixed samples

Epifluorescence images were acquired on an inverted microscope (Nikon Eclipse Ti) with a 20X objective (Nikon Plan Fluor NA=0.5). Confocal microscopy images were acquired on an inverted TCS SP8 confocal microscope (Leica) using a 63X objective. 3D reconstructions were performed using the IMARIS software.

#### Scanning electron microscopy

Cultures were fixed in 2% glutaraldehyde in 0.1 M PBS at pH 7.4. They were dehydrated in a graded series of ethanol solutions and then dried by the CO_2_ critical-point method using EM CPD300 (Leica Microsystems). Samples were mounted on an aluminum stub with a silver lacquer and sputtercoated with a 5 nm platinum layer using EM ACE600 (Leica Microsystems). Acquisitions were performed using a GeminiSEM 500 (Zeiss).

#### Live cell imaging

To stain the nuclei in live cells, Hoechst 33342 (Sigma H3570) diluted 1/5000 in PBS was incubated for 3 min with cells. For visualization of the cell membrane, Cell Mask Orange (ThermoFisher C10045) diluted 1/1000 in culture medium was incubated for 5-10 min with cells. Subsequent live recordings of HUVECs were performed using an automated inverted microscope (Nikon Eclipse Ti) equipped with temperature and CO_2_ regulation and controlled by the NIS software (Nikon). Images were acquired with a 20X objective (Nikon Plan Fluor NA=0.5) for 3 h at 5 min intervals.

#### Atomic force microscopy

For AFM measurements, nuclei were stained with Hoechst 3 h before the beginning of the measurements. Experiments were performed on a NanoWizard 4 BioAFM scanning force microscope (JPK/Bruker) mounted on a Nikon ECLIPSE Ti2-U fluorescent microscope. A probe with a circular symmetric rounded tip with a typical radius of curvature of approximately 30 nm (0.03-0.09 N/m) (uniqprobe qp-BIOAC-CI, from NANOSENSORS^TM^) was used. The spring constant was determined upon calibration by the thermal noise method. Quantitative imaging (QI) (JPK, Berlin, Germany) was conducted in water at 37°C, with a setpoint of 0.8 nN for 90 ms (corresponding to an indentation of approximately 0.5-1 µm). Young’s modulus values were extracted using the Hertz model (Poisson’s ratio of 0.5) and averaged over the nuclear area. Two types of experiments were performed: nuclei in different categories of deformation (based on their shapes and visual classification) were measured once, or single nuclei were measured repeatedly during time (once every hour for a maximum of 7 h).

### Data analysis

#### Analysis and classifications of nuclear shapes

Detection and morphometric analysis of nuclei were performed using a custom-made Matlab code. Briefly, DAPI staining images were used and binarized for subsequent detection of the nuclei. Different morphological parameters were extracted: nuclear (projected) area, nuclear aspect ratio (ratio of major to minor axis length), and measures of tortuosity such as solidity (ratio of area to convex area). From this morphometric analysis, caged nuclei were classified using the following criteria: minor axis < 7 µm and absolute orientation angle < 6° (0° being the orientation of the grooves). Nuclear volume was analyzed independently using the IMARIS software.

#### Analysis of cytoskeletal organization around caged nuclei

Caged nuclei were rotated vertically and cropped from confocal stacks. For each frame in the z direction, a plot profile of the fluorescence intensity across the nucleus minor axis (x axis) of the different cytoskeletal elements was generated and averaged over the entire nuclear length (y axis). The plot profile as a function of depth was then calculated by averaging all grey values for each z and normalizing by the maximum intensity.

#### Analysis of nuclear dynamics

Nuclei were automatically detected and tracked from Hoechst-stained cell recordings using the TrackMate plugin in Fiji to extract whole population cell speed. For more precise analysis of dynamic deformations, caging phases were manually identified on a smaller subset of cells to extract their mean duration and frequency (defined as the number of caging phases for HUVECs or transitions to an adjacent groove for myoblasts, divided by the total time of tracking).

### Data representation and statistical analysis

In all boxplots, the central bar represents the median, the bottom and top edges of the box indicate the 25th and 75th percentiles, respectively, and the whiskers denote the range of minimum to maximum values, excluding outliers. All analyses are based on at least 3 independent experiments. Statistical analyses were performed using the GraphPad Prism software. The Student t-test and the Mann-Whitney t-test were used to compare null hypotheses between two groups for normally and non-normally distributed data, respectively. Multiple groups with a normal distribution were compared by a one-way ANOVA followed by Tukey’s or Fisher’s posthoc test. The number of data points for each experiment, the specific statistical tests, and the significance levels are noted in the corresponding figure legends. On graphs, significance levels are represented with asterisks. In case of multiple conditions, the colors of the asterisks match the condition to which the statistical comparison is made, progressing from left to right.

## Acknowledgements

We thank the MyoLine platform of the Institute of Myology in Paris for providing the myoblast cell lines. We thank P. Mahou and the Polytechnique Bioimaging Facility for assistance in imaging. The Polytechnique Bioimaging Facility is supported by Région Ile-de-France (interDIM) and Agence Nationale de la Recherche (ANR-11-EQPX-0029 Morphoscope2, ANR-10-INBS-04 France BioImaging). This work was supported in part by an endowment in Cardiovascular Bioengineering from the AXA Research Fund (to AIB) and postdoctoral fellowships from the Lefoulon-Delalande Foundation and the Bettencourt-Schueller Foundation (to CL). The present work benefitted from equipment supported by a Leducq Foundation Grant for Research and Technology Platforms.

## Data Availability

The data that support the findings of this study are available from the corresponding author upon reasonable request.

