## Supplementary Figures for "Microgroove substrates unveil topography-driven, dynamic 3D nuclear deformations"

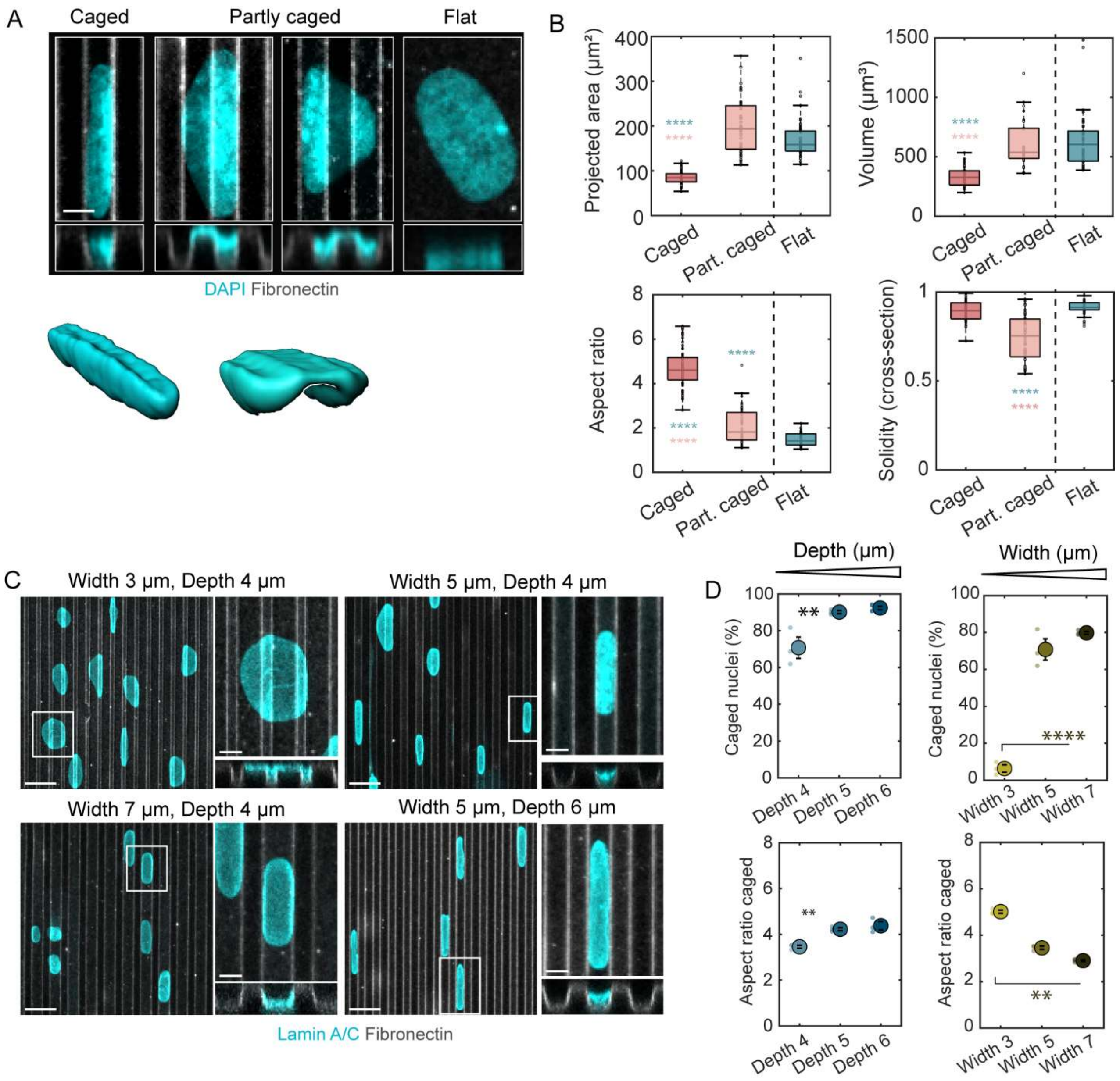


**Figure S1: Characterization of nuclear deformations on microgrooves in myoblasts. (A)** Z-projection images, cross-sections and 3D reconstructions of the different classes of nuclei observed on microgrooves with DAPI. Scale bar 5 µm. **(B)** Morphological characterization of the different classes of nuclei observed on microgrooves and on control flat surfaces. Projected area and aspect ratio were quantified on z-projections images, solidity (tortuosity) on the cross-section images, and volume on 3D reconstructions. n=28-66 cells/category from 3 independent experiments. **(C)** Nuclei (stained for lamin A/C, cyan) on microgrooves (grey) of different dimensions. Scale bars 10 µm, 5 µm (zoom-ins). **(D)** Quantification of the percentage of caged nuclei and aspect ratio of caged nuclei for different groove depths (width = spacing = 5 µm) or groove widths (spacing = 5 µm, depth = 4 µm). Dots represent individual experiments and error bars represent standard error of the mean (SEM). n=3 independent experiments. For all plots: one-way ANOVA, Fisher's post-test (** $p < 0.01$; **** $p < 0.0001$).

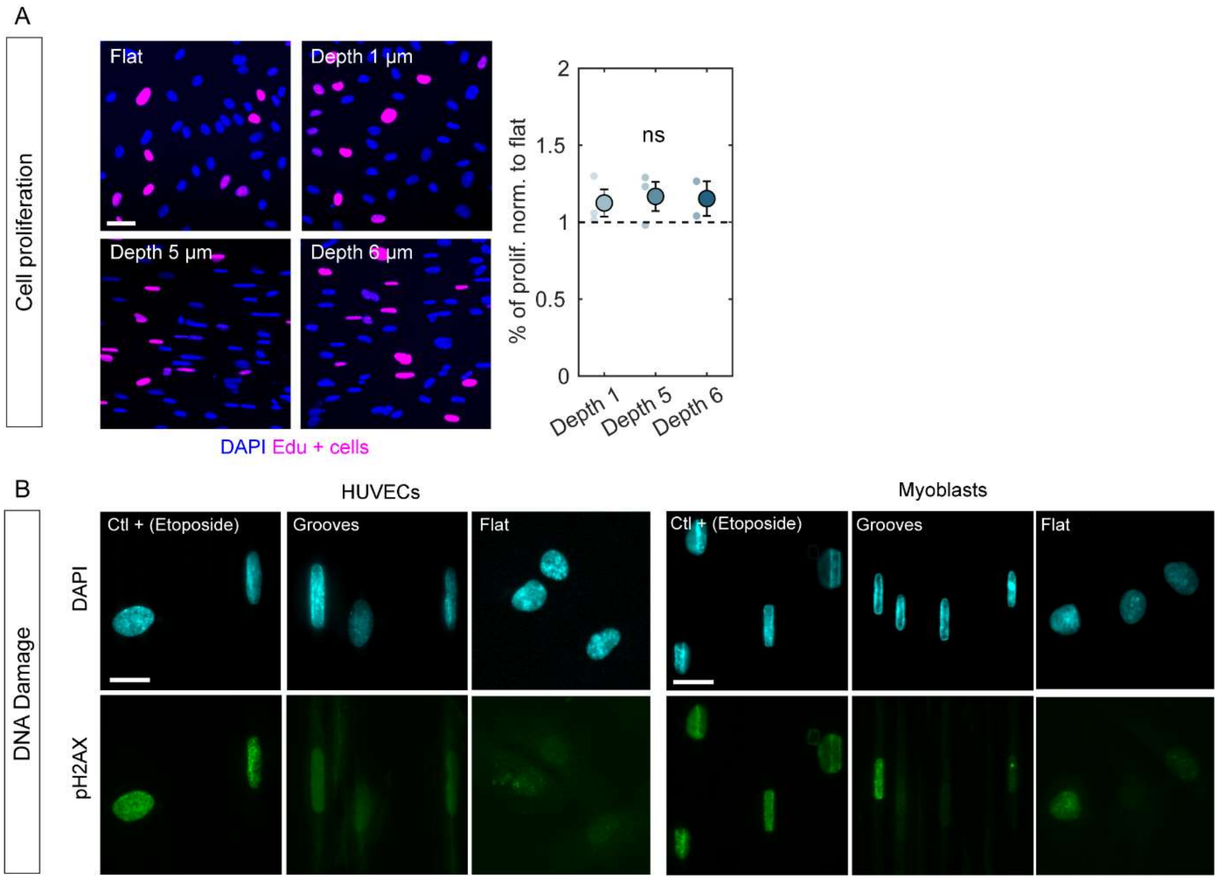


**Figure S2: Functional consequences of nuclear deformations on microgrooves. (A)** EdU assay was used to assess cell proliferation for different groove depths compared to flat PDMS surfaces. Quantification shows the ratio of the percentage of EdU-positive cells on microgrooves to those on flat substrates. **(B)** Immunostaning against phospho-H2AX was used to assess DNA damage in HUVECs or myoblasts cultured on microgrooves or on flat PDMS surfaces. For a positive control, cells were incubated with etoposide (50 mM) for 3 h to induce chemical DNA damage. Scale bars 20 µm.

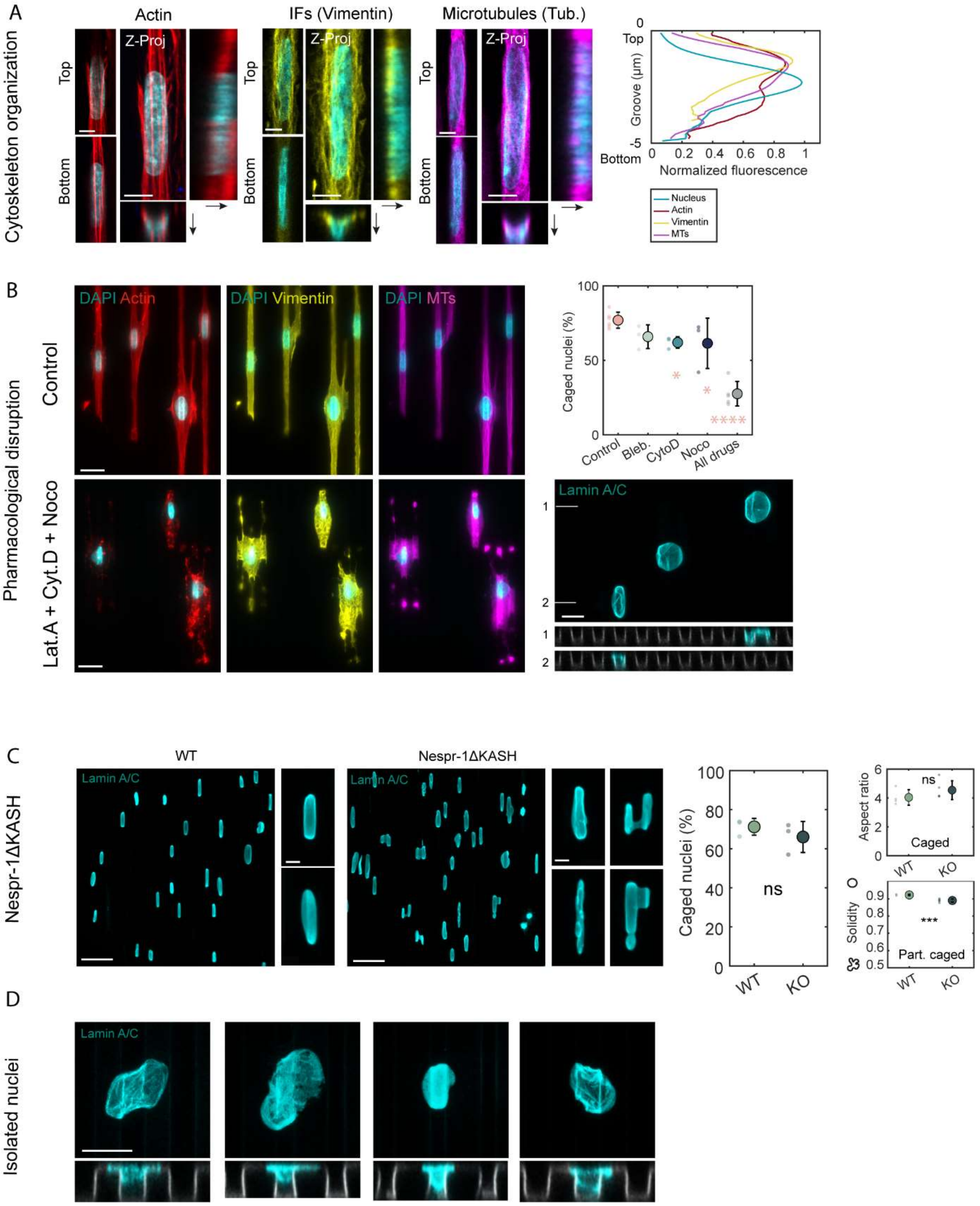


**Figure S3: Organization and role of the cytoskeleton in nuclear deformations on microgrooves in myoblasts. (A) Left:** Z-projections and cross-sections showing the organization of actin (red), intermediate filaments (vimentin, yellow), and microtubules (magenta) around caged nuclei. Scale bars 5 µm. **Right:** Quantification of the normalized fluorescence intensity for the three cytoskeletal networks and the nucleus as a function of depth (from top to bottom of the grooves) for caged nuclei. **(B) Left:** Immunostaining for actin (red), intermediate filaments (vimentin, yellow), and microtubules (magenta) in control cells or cells treated with latrunculin A (Lat.A) + cytochalasin D (Cyto.D) + nocodazole (Noco) on microgrooves (5x5x5 µm, vertical). Scale bar 20 µm. **Right:** Quantification of the percentage of caged nuclei for the different pharmacological treatments: control (DMSO), blebbistatin (Bleb.), cytochalasin D (Cyto.D), nocodazole (Noco), or latrunculin A + cytochalasin D + nocodazole (All drugs). Dots represent individual experiments and error bars represent standard deviations. n=3 to 7 independent experiments. One-way ANOVA, Fisher's post-test (* p < 0.1; **** p < 0.0001). Z-projection and cross-sections of nuclei stained for lamin A/C and treated with latrunculin A + cytochalasin D + nocodazole. Scale bar 10 µm. **(C)** WT or Nesprin1-mutated (Nespr-1ΔKASH) nuclei on microgrooves stained for lamin A/C. Scale bars 50 µm, 5 µm (insets). Quantification of the percentage of caged nuclei, aspect ratio of caged nuclei, and solidity of partly caged nuclei. Dots represent individual experiments and error bars represent standard deviations. n=3 independent experiments. Student t test (* p < 0.1; ** p < 0.01; *** p < 0.001; **** p < 0.001). **(D)** Z-projections and cross-sections of 4 different isolated nuclei deposited on microgrooves and stained for lamin A/C. Scale bar 10 µm.

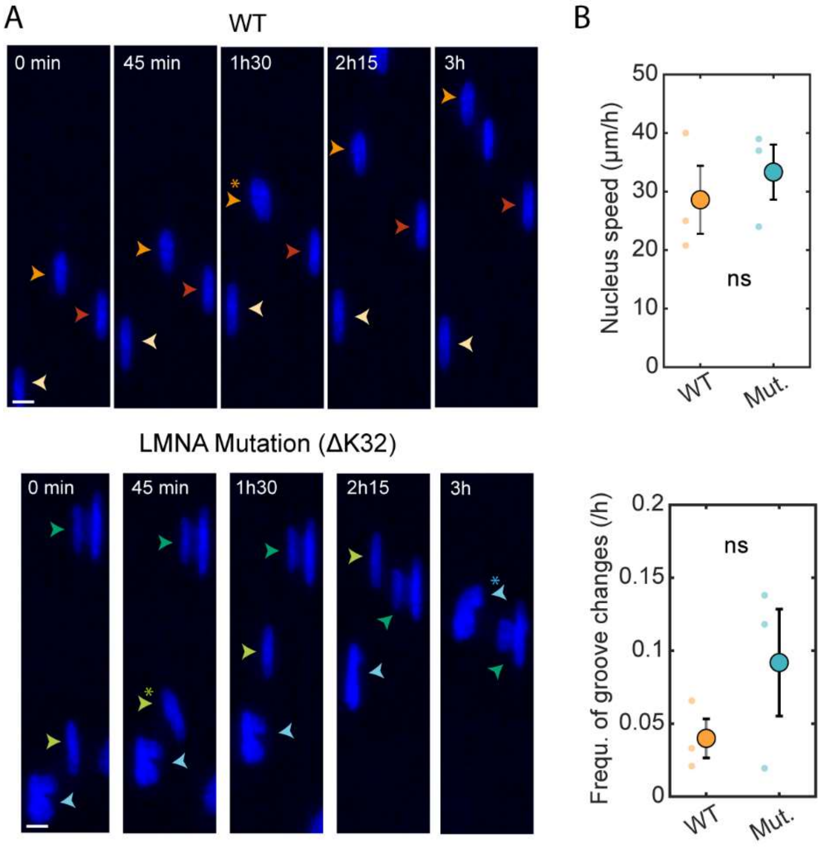


**Figure S4: Dynamics of LMNA mutated myoblasts on microgrooves. (A)** Images extracted from time-lapse recordings of WT or mutated (LMNA mutation ΔK32) myoblast nuclei (stained with Hoechst, blue) on microgrooves (5x5x5 µm, vertical). Arrowheads follow the path of a single nucleus, and stars indicate uncaging phases. Scale bar 10 µm. **(B)** Quantification of the mean nucleus speed and frequency of groove changes. Dots represent individual experiments and error bars represent standard error of the mean (SEM). n=3 independent experiments. Student t test.

### Supplementary movies

**Movie S1:** Time lapse recording of a HUVEC with nuclei stained with Hoechst on microgrooves (5x5x5 µm, vertical). Time interval 15 min, scale bar 20 µm.

**Movie S2:** Time lapse recording of a myoblast with nuclei stained with Hoechst on microgrooves (5x5x5 µm, vertical). Time interval 15 min, scale bar 20 µm.

**Movie S3:** Time lapse recording of a HUVEC with nuclei stained with Hoechst and cell membrane stained with CellMask, on microgrooves (5x5x5 µm, vertical), showing correlated cell and nuclear behavior. Time interval 5 min, scale bar 20 µm.

**Movie S4:** Time lapse recording of a HUVEC with nuclei stained with Hoechst and cell membrane stained with CellMask, on microgrooves (5x5x5 µm, vertical), showing decorrelated cell and nuclear behavior. Time interval 5 min, scale bar 20 µm.
